# Evaluation of Metagenome Binning: Advances and Challenges

**DOI:** 10.1101/2025.02.21.639465

**Authors:** Yazhini Arangasamy, Étienne Morice, Annika Jochheim, Benjamin Lieser, Johannes Söding

## Abstract

**Background:** Several recent deep learning methods for metagenome binning claim improvements in the recovery of high quality metagenome-assembled genomes. These methods differ in their approaches to learn the contig embeddings and to cluster them. Rapid advances in binning require rigorous benchmarking to evaluate the effectiveness of new methods. We have benchmarked newly developed state-of-the-art deep learning binners on CAMI2 datasets, including our own, McDevol.

**Results:** The results show that COMEBin and GenomeFace give the best binning accuracy, although not always the best embedding accuracy. Interestingly, post-binning reassembly consistently improves the quality of low coverage bins. We find that binning coassembled contigs with multi-sample coverage is effective for low coverage dataset while binning multi-sample contigs with multi-sample coverage (‘multi-sample’) is effective for high-coverage samples. In multi-sample binning, splitting the embedding space by sample before clustering showed enhanced performance compared to the standard approach of splitting final clusters by sample.

**Conclusions:** COMEBin and GenomeFace emerged as the top-performing tools overall, with MetaBAT2 and GenomeFace demonstrating superior speed. To facilitate future development, we provide workflows for standardized benchmarking of metagenome binners.

## Introduction

Shotgun metagenomics is widely used to study the genetic diversity, taxonomic composition and metabolic potential of microbial communities. It allows the reconstruction of metagenome-assembled genomes (MAGs), which are valuable for building genomic catalogs, profiling genetic variations, studying accessory genes in strains, and analyzing functional repertoires of microbiomes [1–9]. Since many microbes lack close relatives in reference databases, MAG reconstruction requires an unsupervised approach [10]. Our study focuses on metagenome binning, a key step in reconstructing MAGs without recourse to any reference genomes.

Metagenome binning is the process of clustering contigs assembled from shotgun reads based on their putative genomic origin, resulting in MAGs. The quality of MAGs is often lower than reported values, highlighting the need for further improvements in binning [11, 12]. Metagenome binning clusters the contigs, which are represented as a vector of two types of information: (i) *k* - mer frequencies (typically tetramers), which are similar within the same genome but differ across genomes from different genera. This feature is effective for distinguishing contigs at the genus-level but not at the species or strain level, and (ii) contig coverage in the sample, which is the average number of aligned reads per position, correlating with genome abundance. If multiple samples are available, this information source is very valuable because contigs from the same genome will exhibit similar coverage profiles across samples, providing complementary information for distinguishing contigs at the species level [13]. In addition to these two primary sources, some tools integrate taxonomic annotations [14, 15] and assembly graphs, which represent links between contigs in the de Bruijn graph [16, 17]. For a comprehensive overview, we refer to a recent review [18].

State-of-the-art binners train deep networks to learn lower-dimensional embeddings from the input coverage and *k* -mer frequency vectors of each contig, and perform clustering in the embedding space. For example, the first deep learning binner, VAMB [19], trains a variational autoencoder to learn the embedding (Fig. **1**a). SemiBin [15] introduced contrastive learning to train the network to keep similar contig pairs close together, while dissimilar pairs are placed far apart in the embedding space. To generate contig pairs, data augmentation was used, which involves splitting contigs into shorter fragments and recomputing their coverage and *k*-mer frequencies [14]. SemiBin also provided 11 pre-trained models tailored to specific environments, such as human gut and soil, that can be used to infer the contig embedding space without training. COMEBin [20] extended contrastive learning by using more extended pairs and processing coverage and *k* -mer data separately. In a novel approach, GenomeFace [21] infers the embedding space directly from the input contig vectors using two pre-trained networks. One is trained on *k* mer frequencies with *k* values ranging from 1 to 10 using random contigs from 43,000 curated microbial genomes to predict their taxonomic label from the embeddings. Another network is a transformer model trained on read coverage data using permutation-invariant properties.

**Figure 1:**
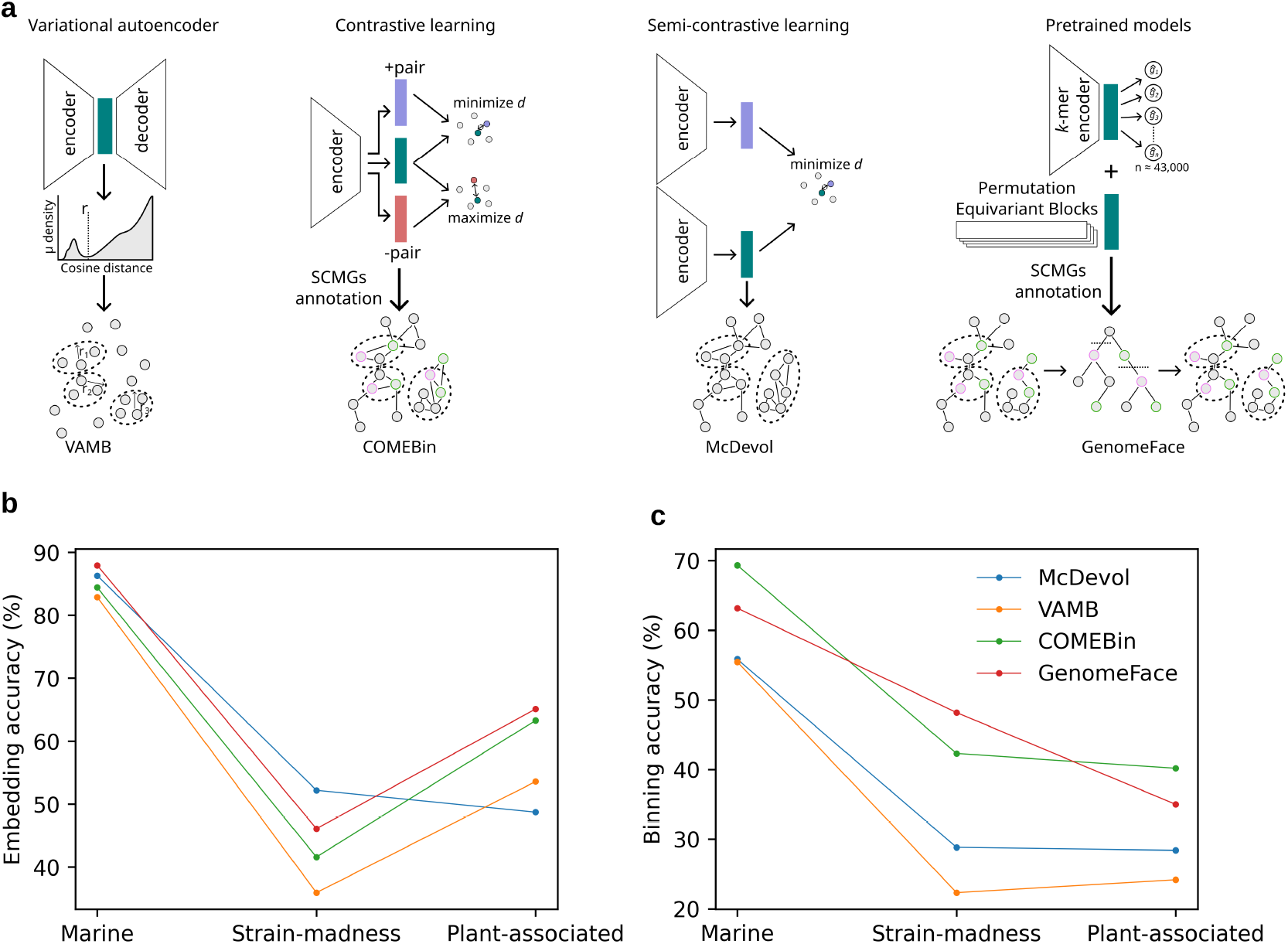
Overview of deep learning binning tools and their accuracy on gold standard coassembled contigs. **a,** Schematic overview of neural network models used in deep learning binning. The symbol *d* denotes the distance between the embedding feature vectors of an augmented contig pair. Blue - positive pair, red - negative pair. **b**, Cross-validated quality of embeddings quantified as the average accuracy of a multi-class linear classifier on the validation datasets. **c**, Accuracy of binning after clustering, as calculated by AMBER [30].

Besides variations in deep learning models, these tools differ in the clustering algorithms used to group contigs in the feature embedding space. VAMB uses iterative density-based clustering, while COMEBin uses the Leiden algorithm and GenomeFace uses hierarchical clustering on a geodesic minimum spanning tree constructed from the embedding space (Fig. **1**a). COMEBin and GenomeFace optimise the clustering process based on measures of completeness and purity derived from the occurrence of single-copy marker genes in the bins [22, 23].

Binning can be performed using one of three approaches: (1) coassembly multi-sample binning, where reads from multiple samples are pooled, coassembled, and binned using abundance across all samples; (2) multi-sample binning, where reads are assembled per sample, contigs are binned collectively using multi-sample abundance, and bins are split by sample origin; and (3) single-sample binning, where contigs are assembled per sample and binned using only their own sample’s abundance [24]. Each approach has trade-offs. Coassembly multi-sample binning increases coverage, enabling the assembly of low-abundance genomes. However, pooling reads from multiple samples introduces higher strain diversity, leading to fragmented assemblies due to ambiguities in contig extension across conserved regions [25]. Single-sample binning is often used in large-scale metagenomics studies due to its computational efficiency [4, 26, 27]. While it reduces strain-diversity-related issues, it does not exploit multi-sample abundance data, poorly recovers low-abundance species, and produces highly similar MAGs across samples, making dereplication essential. A prior study found that sample-wise assembly produces a larger non-redundant gene set than coassembly [25]. VAMB [19] demonstrated that multi-sample binning yields more high-quality, strain-resolved bins than single-sample binning. However, no study has compared the non-redundant results of multi-sample binning with the results of coassembly multi-sample binning.

Rapid advances in metagenome binning and diverse sampling strategies underscore the need for a critical evaluation of state-of-the-art methods to assess their effectiveness in generating high-quality MAGs. In this study, we benchmark recent deep learning-based binning tools using three CAMI2 datasets of varying complexity in genome coverage, contig contiguity and strain diversity. We also include our own binner McDevol, a new deep learning model that employs a semi-contrastive approach to binning. A key challenge in metagenomics datasets is the ambiguity in identifying negative pairs for contrastive learning without taxonomic labels. McDevol’s model addresses this by eliminating the need for negative augmented pairs, allowing more effective learning of embedding representations.

Using gold-standard contigs, we independently evaluated the accuracy of the embedding space prior to clustering and the binning accuracy achieved after clustering. Using MEGAHIT-assembled contigs, we compared the number of bins meeting quality criteria produced by various deep learning methods. In addition, we present the first comparative analysis of metagenomic binning performance after post-binning reassembly and across the described three different binning approaches.

## Results

### Accuracy of feature embedding space and clustering in deep learning binners

Binning tools based on deep learning models typically operate in two phases: (i) learn an embedding in a lower dimensional space for each contig and (ii) clustering contigs represented by their embedding vectors into genomic bins. To compute an embedding function for each contig represented by a feature vector containing its abundances across various samples and its *k*-mer composition, typically across 4-mers, deep neural networks have gained popularity recently. Our first goal was to evaluate the accuracy of contig embeddings generated by different deep learning-based binners, independent of the ensuing clustering process. We evaluated four methods: VAMB (variation autoencoder), COMEBin (contrastive learning), McDevol (semi-contrastive learning) and GenomeFace (pre-trained models). Fig. **1**a summarizes the different learning objectives during training the embedding neural networks. The performance of the binning tools was evaluated by binning gold standard contigs from the CAMI2 marine, strain-madness and plant-associated datasets [28]. The datasets consist of simulated paired-end reads and coassembled contigs, labelled with their source genomes. Contigs were generated by selecting genomic regions from source genomes where simulated reads aligned with a coverage of at least 1 [29]. These datasets exhibit substantial variation in genome fraction covered by the simulated reads (26%–90%), contig contiguity (NGA50 ranging from 87,911 to 682,777 bp), and strain diversity (81–395 strains).

To assess the accuracy of the embeddings, we trained a multi-class linear classifier with softmax output using the embedding feature vectors as input to predict the correct genome, and we evaluated its accuracy through five-fold cross-validation (Methods). Classification accuracy was computed as the ratio of correctly classified contigs to the total, based on ground truth labels. Fig. **1**b reports the embedding accuracy of the four tools in three CAMI2 datasets. The average accuracy was 85.4% for the marine dataset, 44.0% for the strain-madness dataset and 57.7% for the plant-associated dataset, suggesting that the embedding accuracy is strongly influenced by the dataset complexity. For the marine dataset, GenomeFace achieved the highest embedding accuracy (88.0%), followed by McDevol (86.3%), COMEBin (84.4%) and VAMB (82.9%). For the strain-associated dataset, McDevol outperformed the others with an accuracy of 52.2%, ahead of GenomeFace (46.1%), COMEBin (41.6%) and VAMB (35.9%). For the plant-associated dataset, GenomeFace again achieved the highest accuracy (65.1%), followed by COMEBin (63.3%), VAMB (53.6%) and McDevol (48.7%). Overall, GenomeFace was most effective on low coverage datasets (marine and plant-associated), while McDevol performed best on the high genome coverage dataset (strain-madness). McDevol’s poor embedding accuracy for the plant-associated dataset can be attributed to the inherently low read coverage. As shown in Supplementary Fig. 1, the plant-associated dataset has the lowest median total coverage (2.8), compared to marine (3.7) and strain-madness (128.8) datasets. Since McDevol relies on multinomial sampling to generate augmented coverage profiles, low coverage increases the noise in the augmented profiles, thereby reducing the effectiveness of neural network learning.

We evaluated the accuracy of the bins obtained after clustering the embedding space using AMBER [30]. AMBER’s binning accuracy is defined as the average proportion of correctly clustered base pairs into assigned genomic bins (true positives) relative to the total base pairs binned. Fig. **1**c shows the binning accuracy of four binners. The average values for the marine, strain-madness and plant-associated datasets were 60.9%, 35.4% and 31.9%, respectively. This trend reflects the accuracy of the embedding space and highlights the challenges posed by the strain- and plant-associated datasets, particularly due to high fragmentation (Methods). COMEBin achieved the highest binning accuracy for the marine (69.3%) and plant-associated (40.2%) datasets. GenomeFace showed the best binning accuracy for the strain-madness dataset (48.2%). In Supplementary Fig. 2a-c, we compared bin purity and bin completeness. Bin purity is calculated as the proportion of base pairs in a bin that overlap with the assigned genome. Bin completeness is calculated as the proportion of base pairs of the genome covered by the assigned bin. VAMB achieved the best average purity for the marine dataset (95.1%), while GenomeFace achieved the best average purity for the strain-madness (77.9%) and plant-associated (96.5%) datasets. COMEBin achieved the highest average completeness for the marine dataset (64.9%), while McDevol showed the highest completeness for the strain-madness (77.8%) and plant-associated (27.9%) datasets.

On the basis of AMBER’s completeness and contamination (one minus purity), we classified bins meeting the criteria of 90% completeness and 5% contamination as high quality, 70% completeness and 10% contamination as medium quality, and 50% completeness and 10% contamination as low quality. The total number of bins in each category was compared between the binning tools. Supplementary Fig. 2d-f shows that COMEBin generated the highest number of high-quality (312, 47, and 84), medium-quality (399, 62, and 93), and low-quality (454, 70, and 100) bins for the marine, strain-madness, and plant-associated datasets, respectively. GenomeFace was the second best for the plant-associated dataset with 83 high, 99 medium and 102 low-quality bins. Although McDevol achieved comparable embedding accuracy for marine and strain-madness datasets, COMEBin and GenomeFace outperformed it in binning accuracy and produced more high-quality bins. This highlights the effectiveness of COMEBin and GenomeFace in using single-copy marker genes (SCMGs) during clustering to improve the quality of the final bins.

### Benchmarking binning performance on real coassembled contigs

We benchmarked the binning performance of deep learning methods on real metagenomic contigs by coassembling simulated reads pooled across all samples using MEGAHIT [31]. We assessed binners with CheckM2, which predicts the completeness and contamination of each bin through neural network models [32]. We performed binning of these coassembled contigs using read coverage from all metagenomic samples (“coassembly multi-sample”). To compare binning results of deep learning methods with traditional binning approaches, we evaluated the number of high, medium and low-quality bins produced by MetaBAT2, MaxBin2, CONCOCT, and MetaWRAP, a bin-refinement tool. Fig. **2**a reports that the number of bins produced across the three quality categories for the marine dataset were comparable between GenomeFace (108 high-quality, 168 medium-quality, and 199 low-quality bins), MetaBAT2 (110, 167, and 198), and COMEBin (105, 170, and 197) outperforming the other tools. Fig. **2**b shows that for strain-madness dataset, GenomeFace (13 high-quality, 19 medium-quality, and 20 low-quality bins), MetaBAT2 (14, 17, and 19), and McDevol (12, 18, and 19) yielded more quality bins than other tools. However, the differences in the numbers of bins produced by these tools are small. Results in Fig. **2**c indicate that COMEBin (23 high-quality, 49 medium-quality, and 54 low-quality bins), MetaBAT2 (22, 47, and 49), and GenomeFace (17, 41, and 55) generated higher numbers of bins across the three quality categories compared to other tools for the plant-associated dataset. Despite being a traditional binner not using machine learning, MetaBAT2 achieved performance comparable to that of recent deep learning-based binners across all three datasets.

**Figure 2:**
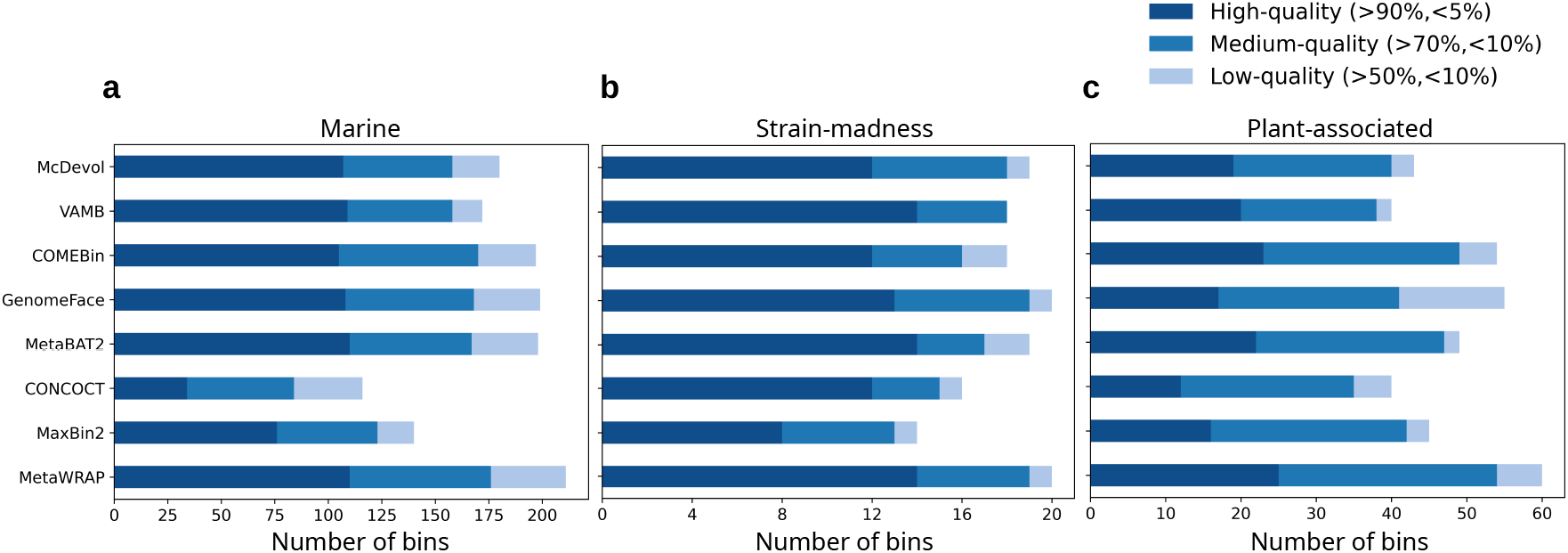
Binning performance on real coassembled contigs with multi-sample coverage. The number of high-quality (dark blue), medium-quality (medium blue) and low-quality (light blue) bins generated for **a)** marine, **b)** strain-madness and **c)** plant-associated datasets as predicted by CheckM2 [32].

The MetaWRAP bin-refinement module, which integrates bins from MetaBAT2, MaxBin2, and CONCOCT, generated the highest total number of bins across the three quality categories. MetaWRAP produced 110 high, 176 medium, and 211 low-quality bins for the marine dataset, and 25, 54, and 60 bins, respectively, for the plant-associated dataset. Compared to the results from the top-performing deep learning binner GenomeFace (199 and 55 bins), the total bin counts produced by MetaWRAP differed by 12 bins for the marine dataset and 5 bins for the plant-associated dataset. For the strain-madness dataset, MetaWRAP produced comparable numbers of bins across the three quality categories (14 high-quality, 19 medium-quality, and 20 low-quality bins) to those produced by the GenomeFace (13, 19, and 20 bins). Compared to the best results from MetaBAT2 among the three tools integrated in the ensemble approach, MetaWRAP achieved a higher percentage gain in bin counts for medium-quality bins (5.4% for marine, 11.8% for strain-madness, and 14.8% for plant-associated) and low-quality bins (6.6% for marine, 5.3% for strain-madness, and 22.5% for plant-associated) than for high-quality bins (0% for marine, 0% for strain-madness, and 13.6% for plant-associated). This suggests that the ensemble approach enhances bin recovery for medium and low quality.

### Reassembly benefits low coverage bins

MetaWRAP has demonstrated that reassembly after metagenome binning can improve bin quality [33]. We evaluated the effect of reassembly, using CheckM2, on genomic bins derived from coassembly multi-sample binning. In Fig. **3**, the top bars correspond to the number of bins in three quality categories without reassembly and the middle bars correspond to the number of bins after reassembly of all bins. On average, reassembly resulted in 16 more high-quality bins in the marine and four more high-quality bins in the plant-associated datasets, but no bins were gained for strain-madness. Marine and plant-associated are low coverage datasets while strain-madness is a high coverage dataset. We investigated whether bin coverage is associated with the improvements in bin quality upon reassembly. Supplementary Fig. 3 indicates that reassembly improved completeness and marginally improved purity only for low-abundance bins, where the mean total read coverage of contigs in the bin is equal to or less than 100. On the other hand, reassembly was ineffective or detrimental to completeness and/or purity of bins with high read coverage (mean bin coverage *>* 100) across all three datasets. Reassembly of only bins with coverage *<* 100 achieves an improvement in the number of bins in the three quality categories (bottom bars of Fig. **3**), a clear improvement over no reassembly of bins (Fig. **3**, top bars).

**Figure 3:**
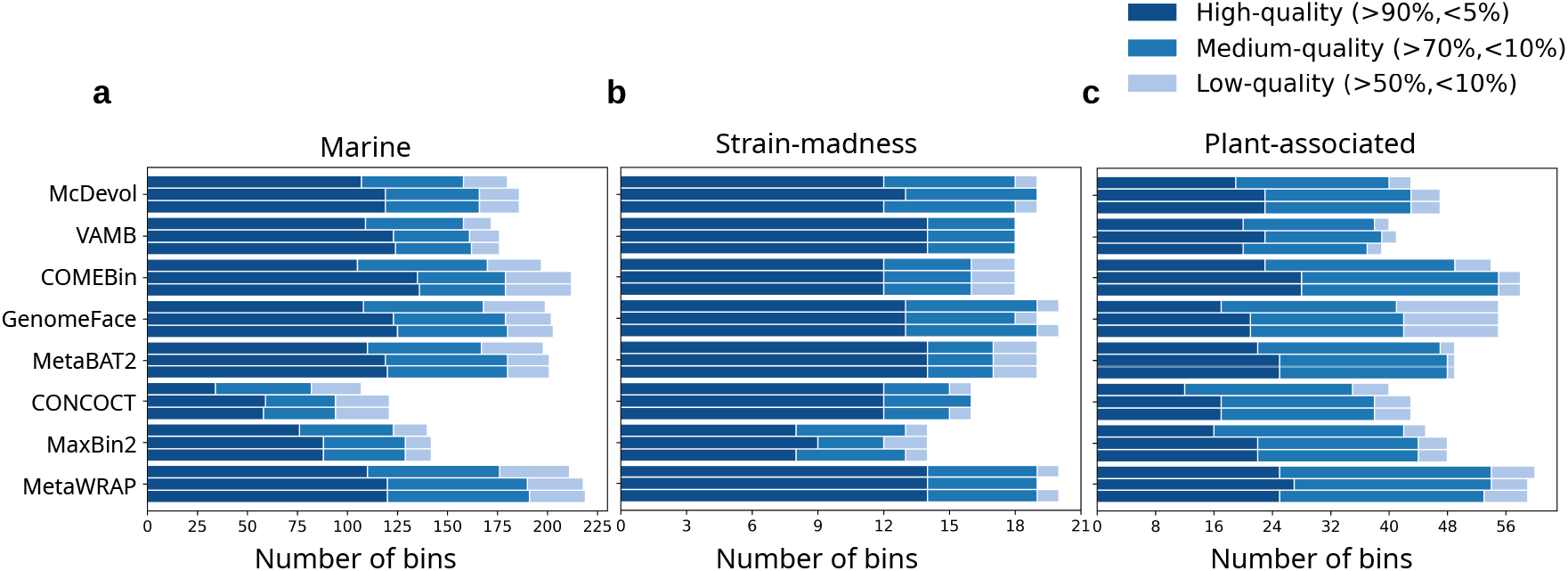
Evaluation of reassembly after binning. The number of bins categorized into high, medium, and low-quality groups for (a) marine, (b) strain-madness, and (c) plant-associated datasets. For each tool, the top bars indicate bin counts before reassembly, the middle bars show bin counts after reassembly, and the bottom bars represent bin counts after reassembling only low coverage bins (mean bin coverage ≤ 100).

Furthermore, improvements by reassembly differed between the three bin quality categories (Supplementary Fig. 4). Reassembly primarily benefited the high-quality bins (mean increase: 22.3% for marine, 22.8% for plant-associated), followed by the medium-quality bins (mean increase: 6.6% for marine, 5% for plant-associated), and had the least positive effect on the low-quality bins (mean: 3.3% for marine, 4% for plant-associated) of all binning tools tested. CONCOCT showed the largest gain with 78.8% and 41.7% additional high quality bins, respectively, in marine and plant-associated datasets, followed by COMEBin with 28.6% more in the marine and MaxBin2 with 37.5% more in the plant-associated datasets.

### Comparison of different binning approaches

As illustrated in Fig. **4**a, Metagenome binning can be performed using different approaches, (1) assembly of pooled reads followed by multi-sample binning (“coassembly multi-sample”), (2) sample-wise assembly followed by multi-sample binning (“multi-sample”), and (3) sample-wise assembly and binning (“single-sample”). In the coassembly multi-sample approach, reads from all metagenomic samples are assembled together and the read coverage of all samples is used to bin contigs. This approach, widely used in metagenome binning, was employed to obtain the results presented in previous sections. The multi-sample approach bins contigs assembled separately for each sample using read coverage data from all samples. The single-sample approach involves both assembly and binning performed independently for each sample. We compared the binning performance of these three approaches for only McDevol, VAMB, COMEBin, GenomeFace and MetaBAT2, all of which had shown competitive results and outperformed other binners in the coassembly multi-sample binning benchmark (Fig. **2**).

**Figure 4:**
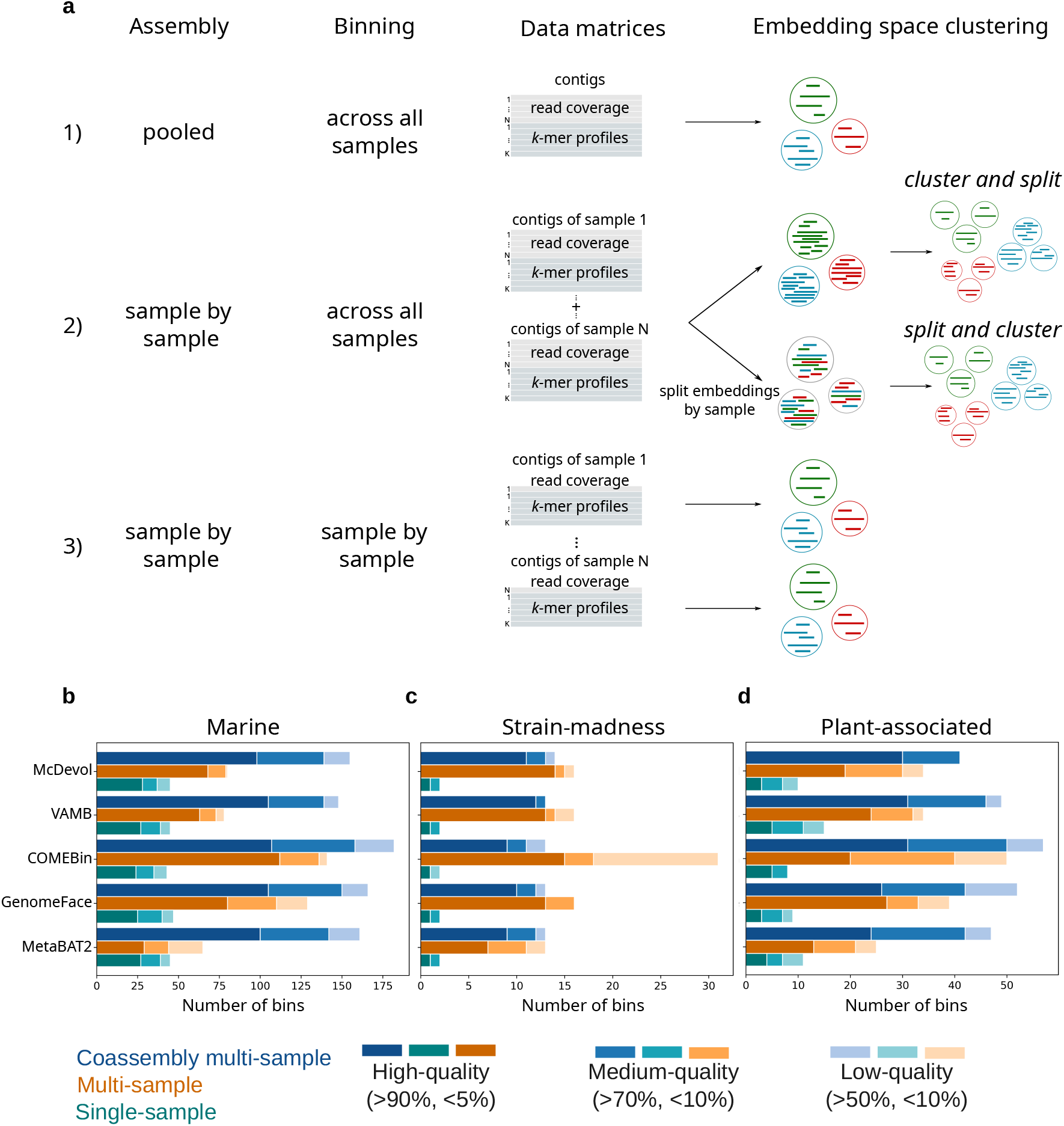
Comparison between different assembly and binning approaches. **a,** 1) Coassembly multi-sample, 2) multi-sample and 3) single-sample. In deep learning methods, multi-sample binning can be performed either by clustering contigs in the embedding space and then splitting clusters by sample (cluster and split), or by splitting contigs in the embedding space by sample and then clustering each sample independently (split and cluster). **b-d**, The number of high, medium and low-quality bins obtained from coassembly multi-sample (blue), multi-sample (orange) and single-sample (cyan) binning for three CAMI2 datasets. The completeness and purity were obtained from the AMBER evaluation [30].

Multi-sample binning can be implemented in two ways: (i) using multi-sample read coverage to bin contigs independently per sample, or (ii) binning contigs from all samples together with multi-sample coverage and then splitting the final bins by sample (Fig. **4**a). In this study, we evaluated the latter strategy as it was applied in previous studies [15, 19] and is resource efficient. In deep learning binners, sample-wise splitting is performed either before clustering (“split and cluster”, as in GenomeFace) or after clustering (“cluster and split”, as in McDevol, VAMB, and COMEBin). We tested these two modes only in McDevol, VAMB and COMEBin, where both modes were possible. We removed redundant bins by selecting the bin with the highest purity per genome across samples (Methods). Supplementary Fig. 5 showed that split and cluster mode consistently outperforms cluster and split mode. For the marine dataset, it produced 72 more high-quality bins in COMEBin and 15 more in McDevol and VAMB, compared to the cluster and split mode. For the strain-madness dataset, COMEBin, VAMB, and McDevol produced 7, 3 and 2 additional highquality bins, respectively. Similarly, for the plant-associated dataset, COMEBin gained 19 more, VAMB gained 9 more, and McDevol gained one more high-quality bins with the split and cluster mode. These results suggest that the split-and-cluster mode is more effective than the cluster-and-split mode for multi-sample binning. Therefore, we used binning results of the split-and-cluster mode to compare with other binning approaches.

Of the three binning approaches compared in Fig. **4**, coassembly multi-sample binning performed best for the marine and plant-associated datasets. It yielded more bins in the three quality categories than multi-sample binning averaged across the five tools, ∼1.8x more for the marine and ∼1.4x more for the plant-associated datasets. However, for the strain-associated dataset, multi-sample binning is advantageous, with ∼1.4x more bins than coassembly multi-sample binning. In particular, COMEBin produced 18 more bins in multi-sample binning than coassembly multi-sample binning.

As expected, multi-sample binning yields significantly more bins than single-sample binning in the three quality categories. COMEBin increased from 43 to 141 bins, GenomeFace from 47 to 129 bins, McDevol from 45 to 80, VAMB from 45 to 78 and MetaBAT2 from 45 to 65 for the marine dataset. A similar trend was observed for the other datasets, with COMEBin showing the largest increase for the strain-madness dataset (2 to 31 bins) and the plant-associated dataset (8 to 50 bins). When considering single-sample binning results alone, cyan bars in Fig. **4**, all binning tools performed similarly, with total bins produced by different tools ranging from 45 to 47 bins for marine, 2 bins for strain madness, and 8 to 15 bins for plant-associated datasets.

### Time and memory usage

We evaluated the run time and peak memory usage of binners for coassembly multi-sample binning of the three CAMI2 datasets. McDevol, GenomeFace, and MetaBAT2 can directly utilize abundance data in tabular format, whereas VAMB and COMEBin require BAM alignment files to extract abundance information, significantly increasing run time. Therefore, we compared run time with and without the preprocessing of alignment files for VAMB and COMEBin. We considered data augmentation as a part of the preprocessing in COMEBin.

Run time comparison in logarithmic scale of minutes in Fig. **5**a reports that MetaBAT2 was the fastest tool for strain-madness and plant-associated datasets, taking approximately 0.2 minutes and 1.6 minutes, 12x and 1.4x faster respectively than the second fastest tool, GenomeFace. For the marine dataset, GenomeFace was the fastest tool, requiring 2.6 minutes, which is 2x faster than MetaBAT2 (5.6 minutes). Unlike other tools, GenomeFace’s run time remained stable regardless of the number of contigs or samples. However, it required an additional 60, 6.6 and 39 minutes on a single CPU node to identify SCMGs in the marine, strain-madness and plant-associated datasets, respectively. McDevol’s run time was approximately twice that of VAMB due to the longer training epochs and the use of a contrastive network model that optimises a larger number of parameters. VAMB’s binning run time was shorter without preprocessing for the strain-madness dataset, suggesting that the overall run time of VAMB is largely due to the preprocessing of multiple BAM files. COMEBin was the slowest tool due to its data augmentation, extended training time, and multiple rounds of Leiden community detection to optimise clusters based on CheckM quality measures.

**Figure 5:**
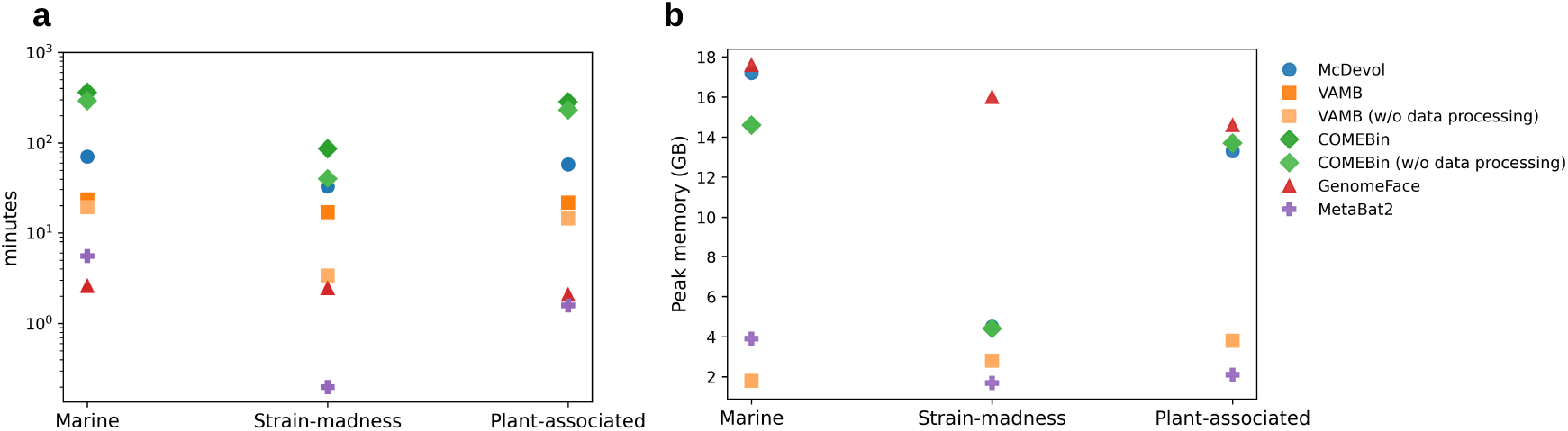
Computational efficiency. **a,** Run time (in minutes) required by state-of-the-art tools for binning the three CAMI2 datasets. The y-axis is plotted on a logarithmic scale. **b**, Peak memory usage (in gigabytes) observed during the execution of each binning tool.

In Fig. **5**b, GenomeFace had the highest peak memory usage with 17.6GB, 16GB and 14.6GB for the marine, strain-madness and plant-associated datasets respectively. McDevol followed with peak memory usage of 17.2GB, 4.5GB and 13.3GB, while COMEBin’s peak memory usage was 14.6GB, 4.4GB and 13.7GB for the respective datasets. VAMB had the lowest peak memory usage of 1.8GB for the marine dataset. MetaBAT2 showed the lowest peak memory usage of 1.7GB and 2.1GB for the strain-madness and plant-associated datasets respectively, which was twice as low as VAMB.

## Discussion and Outlook

Metagenome binning is a challenging clustering problem that has seen considerable progress with advances in representation learning. In this study, we benchmarked tools for metagenome binning. Our work differs from the CAMI2 challenge [28] in four key aspects: (i) we compared recently developed state-of-the-art deep learning binners, (ii) we independently assessed the accuracy of the learned embeddings and the clustering, (iii) we investigated the impact of reassembly after binning, and (iv) we evaluated the performance of co-assembly multi-sample, multi-sample and single-sample binning approaches. Among the methods tested, COMEBin and GenomeFace demonstrated the highest binning performance, surpassing the best results of a single binner in the CAMI2 challenge. Consistent with previous findings [28, 33], the ensemble method MetaWRAP yielded the highest total number of bins; however, our analysis indicates that its benefits are mainly limited to medium- and low-quality bins (Fig. **2**).

Reassembly has been proposed as a post-binning refinement approach to improve MAGs recovery [33–35]. Our results indicate that the benefit of reassembly depends on the read coverage of the bins. It significantly improves the completeness and marginally improves purity for low-coverage bins, while providing no benefit or detriment for high-coverage bins in all three datasets (Fig. **3**). This may be due challenges in initial coassembly. To decide which paths through the de-Bruijn graph to extract as contigs, assemblers try to find the paths with high and consistent coverage. Paths/contigs with low coverage are much harder to distinguish from other paths/contigs with similar, low coverage. During reassembly, only a small subset of reads is assembled, resulting in much simpler the de-Bruijn graphs with fewer ambiguities. In contrast, high coverage reads are already well assembled during the initial co-assembly, making reassembly less beneficial in such cases. Therefore, the decision to use reassembly should be guided by bin coverage. Interestingly, reassembly contributes more to high-quality bins than to medium and low quality bins (Supplementary Fig. 4). We find that the available reassembly module from MetaWRAP has limitations: it doesn’t support interleaved reads, and doesn’t handle reads from multiple samples efficiently, which can lead to memory constraints. To address these issues, we provide a simple reassembly workflow that bypasses MetaWRAP’s read mapping step for reassembly (Supplementary Fig. 9). Our workflow also parallelizes reassembly across bins, improving time efficiency.

Mattock and Watson [24] compared multi-sample binning with single-sample binning and showed significant differences in bin yield. However, this study did not include the widely used coassembly multi-sample binning approach. We performed an extensive comparison of three different binning approaches and report that, given the multi-sample coverage data, the binning of coassembled contigs outperformed multi-sample contigs for the low coverage samples (marine and plant-associated, Fig. **4**). In the marine dataset, coassembly produces longer contigs, resulting in more reliable read coverage and *k*-mer profiles and better binning results (Supplementary Fig. 6). For plant-associated datasets, where both approaches generate contigs of similar lengths, coassembly multi-sample binning remains more effective. We speculate that this is due to the same genomic region being represented by several contig from different samples. When reads are mapped to a concatenated set of contigs, they are assigned to only one of these contigs, resulting in reduced read counts. In contrast, in coassembly multi-sample binning, genomic regions are likely represented by a single contig, allowing unambiguous read mapping. This results in higher counts, more accurate abundance profiles and thus better binning performance.

In the strain-madness dataset, contigs in coassembly multi-sample binning are shorter than contigs in multi-sample binning due to high fragmentation in the coassembly. Current genome assemblers struggle to expand assemblies when reads from other samples belonging to different genomes share highly similar regions, which is common in the strain-madness dataset. By assembling reads of each sample independently, multi-sample binning avoids fragmentation due to ambiguities in possible extensions. Longer contigs have more read counts and less noisy abundance profiles for binning. Thus, multi-sample binning is advantageous for datasets with high strain complexity. However, this advantage is likely contingent on high per-sample coverage, as simulated in the strain-madness dataset. We recommend the “split and cluster” mode, which improves binning results compared to the default “cluster and split” mode in existing multi-sample binning tools (Supplementary Fig. 5). Additionally, redundancy filtering or selecting the best bin across samples may discard valuable genomic information present in unselected bins from other samples. Developing an efficient method to merge bins across samples could further enhance the performance of multi-sample binning.

Runtime evaluation showed that MetaBAT2 and GenomeFace were the fastest tools. COMEBin was the slowest tool, taking up to 2 days for multi-sample binning and causing out-of-memory problems during its clustering step. Addressing these limitations could make COMEBin more suitable for large-scale metagenomic studies. McDevol, a semi-contrastive model, is faster than COMEBin. It showed comparable embedding accuracy to contrastive learning for marine and strain-madness datasets, demonstrating its efficiency in achieving useful embedding space without negative pairs. We find that McDevol is powerful when the metagenomic samples have a high coverage, but it is inferior to other deep-learning binners for samples with many low-coverage genomes. In terms of memory usage, although GenomeFace has the highest peak memory requirement (Fig. **5**b), it remains within the RAM limits of modern computers.

A limitation of this benchmarking is its focus on prokaryotic genomes only, as metagenome binning tools and evaluation methods are standardised for prokaryotes (bacteria and archaea). For example, SCMGs used in supervised clustering for binning, and training datasets for evaluation tools such as CheckM2 [32] and DeepCheck [36], contain predominantly prokaryotic and archaeal genomes. However, TaxVAMB [37] demonstrated improvements in binning both prokaryotic and fungal genomes by incorporating comprehensive taxonomic data. In addition, genome foundation models such as DNABERT-S [38] have improved the accuracy of sequence composition embeddings. With rapid taxonomic annotation tools such as MMseqs2 taxonomy [39], Metabuli [40] and Taxometer [41], future binners combining genome foundation models with taxonomic data in semi- or self-supervised multimodal approaches could improve genome binning across diverse domains of life.

In summary, our work demonstrates that although the yield of MAGs is largely influenced by the complexity of the dataset, the choice of appropriate assembly and binning approaches can significantly affect MAG recovery. For low-coverage datasets, we recommend coassembly multi-sample binning (using tools such as GenomeFace, COMEBin, MetaBAT2, or an ensemble approach) followed by reassembly refinement. For high coverage datasets, multi-sample binning is most effective.

## Methods

### McDevol semi-contrastive deep learning metagenome binner

We developed a new metagenomic binner, McDevol, which follows the BYOL (Bootstrap Your Own Latent) semi-contrastive learning [42] to obtain feature embeddings of contigs from read coverage and *k*-mer frequency profiles. McDevol performs binning in a self-supervised manner and does not require taxonomy assignment or annotation with single-copy marker genes.

#### Inputs and preprocessing

McDevol requires contig sequences and the abundance matrix in tab-separated format. It also accepts raw read contig alignment files in SAM format directly, without the need for time and memory consuming sorting and/or indexing of alignment files. Our alignment parser filters read alignments by sequence identity (≥97%) and read alignment coverage (≥99%). During the calculation of sequence identity, low quality nucleotide positions in the reads (phred score *<* 20) are ignored. If a read aligns equally well to multiple contigs, we distribute the read alignment counts evenly across the aligned contigs. Contig read counts are converted to coverage by multiplying by read length and dividing by contig length. The read coverage values of contigs per sample are then converted into an abundance matrix with which the neural network is trained.

#### Data augmentation and training

The BYOL model trains on only positive augmented pairs (Fig. **1**a). They are created by randomly splitting each contig into smaller fragments, and their read coverage and *k*-mer profiles are independently generated using multinomial sampling and GenomeFace’s composition models, respectively (Supplementary Materials). During training, the input vectors of the augmented pairs are fed into the online and target networks of the BYOL model. The target network is only updated at each training step with a slow moving average of the online network. McDevol trains for 400 epochs using the Stochastic Gradient Descent (SGD) optimiser with a learning rate of 0.1 and incorporates early stopping. The learning rate follows a linear warm-up from 0 to 0.1 over the first 20 epochs, after which it is adjusted using a cosine annealing scheduler. Early stopping is triggered when the validation loss does not improve beyond a small threshold (1*e*^*−*6^) for 10 consecutive epochs. A detailed model architecture is available in the supplementary material.

#### Clustering

To group contigs into bins based on their feature embedding representations, McDevol employs the Leiden community detection algorithm [43]. A nearest-neighbor graph of contigs is constructed using the hnswlib package [44], with default parameters and L2 distance as the similarity metric. This graph is subsequently processed by the Leiden algorithm, utilizing the RBConfigurationVertexPartition optimizer [45] for community detection. The resolution parameter and the minimum number of edges required for community detection are set to their default values of 1.0 and 100, respectively. To ensure biologically meaningful bins, McDevol filters the detected communities to include only those with a minimum total base pair size of 200 kbp.

### CAMI2 benchmark datasets and assembly

We used the gold standard datasets from the CAMI2 assessment study [28], namely marine, strain-madness and plant-associated samples. These datasets are recent, have different strain diversity and are not widely used in the development of binning tools. The marine dataset contains 777 genomes (474 unique and 303 strains) with simulated reads for 10 samples. The strain-madness dataset contains 408 genomes (13 unique and 395 strains) with simulated reads for 100 samples. The plant-associated dataset contains 495 genomes (414 unique and 81 strains) with simulated reads for 21 samples. The gold standard coassembled contigs provided by the CAMI2 study show greater variability in genome coverage and sequence contiguity across the datasets. For marine, the contigs show moderate genome fraction coverage (76.9%) but are contiguous (NGA50 of 682,777 bp). The contigs of the strain-madness set show 90.0% genome fraction coverage but low contiguity (NGA50 of 155,980 bp). The plant-associated contigs have the lowest genome fraction coverage (29.6%) and are more fragmented (NGA50 of 87,911 bp) than the other two contig sets.

We found that the simulated reads had an error rate of 3%, significantly higher than the typical error rate of 0.1 to 1% for standard Illumina short reads [46]. To address this, we used CoCo (git@github.com:soedinglab/CoCo.git), an error correction tool that corrects reads based on *k* - mer counts. Since *k*-mer counts from pooled samples provide more accurate information than single sample data, we corrected reads by counting *k*-mers across all samples together. This error correction reduced the error rate from 3% to 1%. Assembly was performed with MEGAHIT (v1.2.9) [31] using the `--presets meta-sensitive` option for coassembly. For individual sample assemblies, corrected reads were separated by sample and independently assembled. We evaluated the assembled contigs by MetaQuast [47] and found that they cover 44.99%, 8.4% and 14.1% fractions of the genomes for the marine, strain-madness and plant-associated datasets, respectively, while contigs from sample-wise assembly cover 41.1%, 64.7% and 8.8%. The mean contig lengths of the coassembly are 1076bp, 670bp and 954bp, while the same for the multi-sample contigs are 943bp, 1216bp and 1034bp for the three datasets, respectively (Supplementary Fig. 6).

### Metagenome binning benchmark

To perform binning, we generated an abundance matrix by mapping reads to assembled contigs. Strobealign [48] was used to estimate abundance values using the `--aemb` option and to align reads from each sample to contigs to obtain alignment SAM files. McDevol, GenomeFace and MetaBAT2 accept the abundance matrix directly, while VAMB and COMEBin require sorted BAM files as input. We consolidated the abundance values from each sample into a matrix format (similar to MetaBAT2’s *jgi summarize bam contig depths* output) using an in-house Python script. Samtools (v1.19) was used to create sorted BAM files from the alignments. For all binners, only contigs of at least 1000 bp were used for binning, except for MetaBAT2, which showed better performance for contigs above 1500 bp. We ran MetaWRAP for MetaBAT2, CONCOCT, and MaxBin2 tools with assembled contigs and sequencing reads as input. By default, MetaWRAP performed read mapping and generated abundance matrices to facilitate binning. The results from all three tools were then integrated using MetaWRAP’s bin refinement module [33]. The commands used for the benchmarking runs are available in the supplementary material.

### Single-sample and multi-sample binning

For single-sample assembly binning, three samples were randomly selected, assembled and independently binned. The best binning results from these three samples were selected for the comparison. For multi-sample binning, each sample was assembled independently using MEGAHIT [31], but all assembled contigs were binned together. We performed multi-sample binning in cluster-and-split as well as split-and-cluster modes and evaluated results from McDevol, VAMB and COMEBin, as only these tools allowed the evaluation of two modes. Multi-sample binning of the same environmental samples often results in redundant bins due to the presence of the same genomes in multiple samples. To remove redundancy, we mapped each sample’s reads to the contigs using Strobealign [48]. We assigned each contig to the genome that had the highest fraction of its total genomic reads mapped to that contig. Using the purity and completeness measures estimated by AMBER [30], we selected the best bin for each genome among the samples for comparison. The same protocol was followed for coassembly multi-sample and single-sample binning, to compare assessment results with multi-sample binning.

### Evaluation

To evaluate the accuracy of the embedding spaces generated by the binning tools, we used a linear multi-class classifier with softmax output implemented in PyTorch (v1.12.1), each genome being represented by one class. The classifier consists of a single linear layer with the following parameters: ‘learning rate’: 0.001, ‘optimiser’: Adam, ‘epoch’: 300 and ‘Loss’: Cross entropy. For the evaluation, we used short gold-standard pooled contigs and corresponding genome labels for the marine, strain-madness and plant-associated datasets from the CAMI2 study [28]. Each dataset was randomly divided into five groups for 5-fold cross-validation using scikit-learn KFold [49]. We computed the classification accuracy as the ratio of correctly labeled contigs to the total number of contigs in the test dataset. The embedding space accuracy of the binning tool was defined as the average classification accuracy across the test datasets following 5-fold cross-validation. This accuracy was then compared across four different binning tools.

We used AMBER [30] to evaluate the accuracy of metagenome binning and the quality of individual bins. Accuracy of binning is defined as the average of the number of base pairs that overlap with the genome assigned of the bin (TP) over the number of base pairs in the entire dataset, including base pairs not overlapping with the assigned genome (FP) and unassigned base pairs (TP+FP+Unassigned). We computed the average completeness (bp) and average purity (bp) of bins using AMBER [30]. Briefly, average purity (bp) is the fraction of correctly assigned base pairs in each bin, defined as the proportion of base pairs in a bin that belong to the predominant genome in that bin, averaged across all predicted bins generated by the binner. Average contamination is computed as 1 minus average purity. Average completeness (bp) refers to the fraction of base pairs from a genome that are covered by contigs within a given bin, averaged across the completeness values of all genomes in the sample. For bins generated from contigs assembled using MEGAHIT [31], we used CheckM2 [32] to predict completeness and contamination. Bins were classified into three groups based on these two measures: high quality (≥ 90% completeness and *<*5% contamination), medium quality (≥70% completeness and *<* 10% contamination), and low quality (≥ 50% completeness and *<*10% contamination). In all cases, bins with a total length of at least 200kb were selected for comparison.

### Post-binning reassembly

We used the SPAdes assembler (v4.0.0) [50] for reassembly with the `--careful` and `--trusted-contigs` flags. For each bin, reads were extracted from all samples using the ‘extrac-treads’ C++ executable with unsorted sam or bam input files. Reassembly was performed using the bin-specific reads together with the contigs belonging to that bin. Final scaffolds were filtered by a minimum length of 500 bp, and CheckM2 was used to assess the quality of the reassembled bins.

### Reproducibility

We evaluated the run time and peak memory usage of the binning tools on a Cascade Lake 6242 machine equipped with an NVIDIA Quadro RTX 5000 GPU, using 1 CPU with 24 cores and 128 GB of main memory. To ensure reproducibility, we provide Snakemake workflows for coassembly-multi-sample and multi-sample binning. These workflows cover the entire pipeline, including assembly, read mapping, alignment sorting, abundance matrix generation, binning and evaluation (Supplementary Figs. 7 and 8). We have also developed a reassembly workflow that includes read recruitment for each bin, reassembly, scaffold filtering and quality assessment using CheckM2 (Supplementary Fig. 9).

## Declarations

### Ethics approval and consent to participate

Not applicable

### Consent for publication

Not applicable

### Availability of data and materials

Software, scripts used for data analysis, and summary results are available as open source at https://github.com/soedinglab/binning_benchmarking.git. Command lines for running the binning tools and parameters are given in the supplementary material. McDevol’s Python source code is publicly available at https://github.com/soedinglab/McDevol.git along with the compilation and execution instructions. The datasets used from the CAMI2 study are available at https://frl.publisso.de/data/frl:6425521.

## Authors’ contributions

YA and JS conceived of the project. YA and JS designed the algorithm. YA ran benchmarks and generated figures. EM and BL advised on the implementation of McDevol and workflows. AJ advised on pre-processing metagenomic data. YA drafted the manuscript and JS revised it. All authors read and approved the final manuscript.

## Acknowledgements

This work used the Scientific Compute Cluster at GWDG, the joint data center of the Max Planck Society for the Advancement of Science (MPG) and the University of Göttingen. The authors thank Dr Ruoshi Zhang for the critical reading of the manuscript.

