## Supplementary_Information.pdf for "Evaluation of Metagenome Binning: Advances and Challenges"

#### McDevol – Generating feature profiles for positive augmented pairs

We generate augmented pairs by randomly splitting contigs into shorter fragments, each with a minimum length of 1000 bp, and independently generating their read coverage and k-mer profiles. To represent the k-mer features of these augmented pairs, we use the GenomeFace's composition model to generate k-mer feature embedding vectors. For the read coverage profiles, we apply multinomial sampling. Specifically, given the read coverage profiles of the original contigs, we compute the read count profiles, $\vec{x}$ as

$$\text{read count} = \frac{\text{read coverage} \times \text{contig length}}{\text{read length}}$$

The read length is set to 250 for paired-end metagenomic reads of 150 bp. The count profiles are used in Dirichlet sampling to derive probability vectors $p(\vec{\theta}|\vec{x})$, with the concentration parameter vectors $\vec{\alpha}$ set to value 1. These probability vectors are scaled by multiplying them with the total read counts of the fragmented contig. This is approximated as the total read counts of the original contig scaled by the fraction of the length covered by the fragmentation ($X'_{total}$). Using these scaled probabilities $p(\vec{\theta}|\vec{x})_{\text{scaled}}$ as rate parameters, we apply Poisson sampling to generate augmented read count profiles, $\vec{x}'$.

$$p(\vec{\theta}|\vec{x}) = \text{Dir}(\vec{\theta}|\vec{x} + \vec{\alpha})$$

$$p(\vec{\theta}|\vec{x})_{\text{scaled}} \approx p(\vec{\theta}|\vec{x}) * X'_{\text{total}}$$

$$\vec{x}' \approx \text{Pois}(p(\vec{\theta}|\vec{x})_{\text{scaled}})$$

We use Poisson instead of multinomial sampling to efficiently vectorize the count sampling in each training batch. To ensure numerical stability, we iterate the Poisson sampling until no count profile has a total count of zero. We also clamp the count with the maximum observed read count in the batch. We convert the augmented count profiles into read coverage profiles. During neural network training, this procedure is applied twice to generate two augmented read coverage profiles for each contig in the batch. The augmented read coverage profiles are combined with GenomeFace's k-mer embedding vectors to feed into McDevol model.

#### McDevol architecture

McDevol follows the framework of BYOL (Bootstrap Your Own Latent) [1] to learn feature embedding representations ($y_\theta$) from coverage and k-mer inputs. This self-supervised approach involves two networks: an online network and a target network, each designed to process augmented input pairs in parallel. The online network consists of an encoder ($f_\theta$), a projector ($g_\theta$) and a predictor ($q_\theta$). The encoder consists of two fully connected hidden linear layers with an output size of 1024 and a final linear layer that outputs a 512-dimensional embedding vector ($y_\theta$). The output of the hidden layers undergoes batch normalisation, leaky rectified linear unit (ReLU) activation and dropout (rate of 0.1). The encoder output is fed to the projector, which reduces the dimensionality to a 256-dimensional space using a multilayer perceptron (MLP) ($g_\theta$). The MLP consists of a linear layer with an output size of 2048, batch normalisation and leaky ReLU activation, and an output linear layer with a projection size of 256. The predictor ($q_\theta$) uses the same architecture as the projector. The target network consists of encoder and projector with the same architecture as the online network, but gets the model weights from an exponential moving average of the online network parameters with a decay rate of 0.99. The training objective is to minimize the mean squared error, denoted as $L_{\theta,\xi}$, between the normalised vectors from the online predictor ($q_\theta(z_\theta)$) and the target projector ($z'_\xi$). The loss is symmetrized by adding the mean squared error computed on the swapped augmented pair fed to the online and target networks ($\widetilde{L}_{\theta,\xi}$). These are defined as

$$L_{\theta,\xi} = 2 - 2\frac{\langle q_\theta(z_\theta), z'_\xi \rangle}{\|q_\theta(z_\theta)\|_2 \cdot \|z'_\xi\|_2}$$

and

$$\widetilde{L}_{\theta,\xi} = 2 - 2\frac{\langle q_\theta(z'_\theta), z_\xi \rangle}{\|q_\theta(z'_\theta)\|_2 \cdot \|z_\xi\|_2}$$

where $q_\theta(z_\theta)$ and $z'_\xi$ represent the online predictor and target projector outputs, respectively, of an augmented pair. $q_\theta(z'_\theta)$ and $z_\xi$ represent the online predictor and target projector outputs, respectively, of swapped inputs of the same augmented pair. The total loss function is then given by

$$Loss = L_{\theta,\xi} + \widetilde{L}_{\theta,\xi}$$

The model is trained using the stochastic gradient descent optimiser with a batch size of 4096. McDevol was implemented in Python (v3.9) using PyTorch (v1.10.0) and CUDA (v11.0) is enabled when running on a machine with GPUs.

#### Commands used to run metagenome binning tools

Runs correspond to metagenome binning of coassembled contigs (single_pooled_final.contig.fa) using McDevol, VAMB, COMEBin, GenomeFace, MetaBAT2 and MetaWRAP binning modules (MetaBAT2, MaxBin2 and CONCOCT).

```
/usr/bin/time -v mcdevol/mcdevol.py -c sorted_pooled_final.contigs.fa -o
↪  mcdevol_results -a abundances_gf_sorted.tsv --abundformat metabat -n 24

/usr/bin/time -v vamb --outdir vamb_results --fasta
↪  single_pooled_final.contigs.fa --bamfiles bamfiles/*.bam -m 1000 --minfasta
↪  200000 --cuda -p 24

/usr/bin/time -v run_comebin.sh -a single_pooled_final.contigs.fa -o
↪  comebin_results -p bamfiles -t 24

/usr/bin/time -v genomeface -i sorted_pooled_final.contigs.fa -o
↪  genomeface_results -a abundances_gf_sorted.tsv -g marker_hits -m 1000

/usr/bin/time -v metabat2 -i sorted_pooled_final.contigs.fa -a
↪  abundances_gf_sorted.tsv -o /metabat2_results/metabat2_results -t 24 -m 1500

/usr/bin/time -v metawrap binning -o metawrap -t 24 -a
↪  sorted_pooled_final.contigs.fa --metabat2 --maxbin2 --concoct --interleaved
↪  fastqfiles/coco_corrected/*.fastq

/usr/bin/time -v metawrap bin_refinement -o metawrap/bin_refined -t 24 -A
↪  metawrap/metabat2_bins/ -B metawrap/maxbin2_bins/ -C metawrap/concoct_bins/
↪  -c 50 -x 10
```

### 1 Supplementary Figures

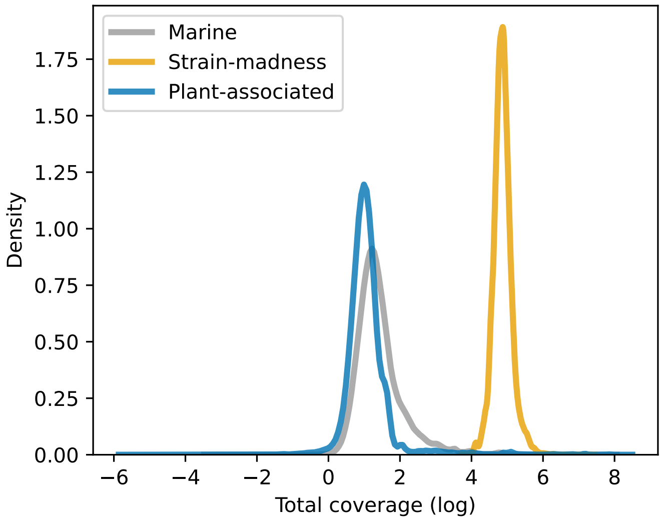


Figure 1: Total read coverage (in log-scale) computed for gold-standard contigs from the marine (grey), strain-madness (orange) and plant-associated (blue) datasets.

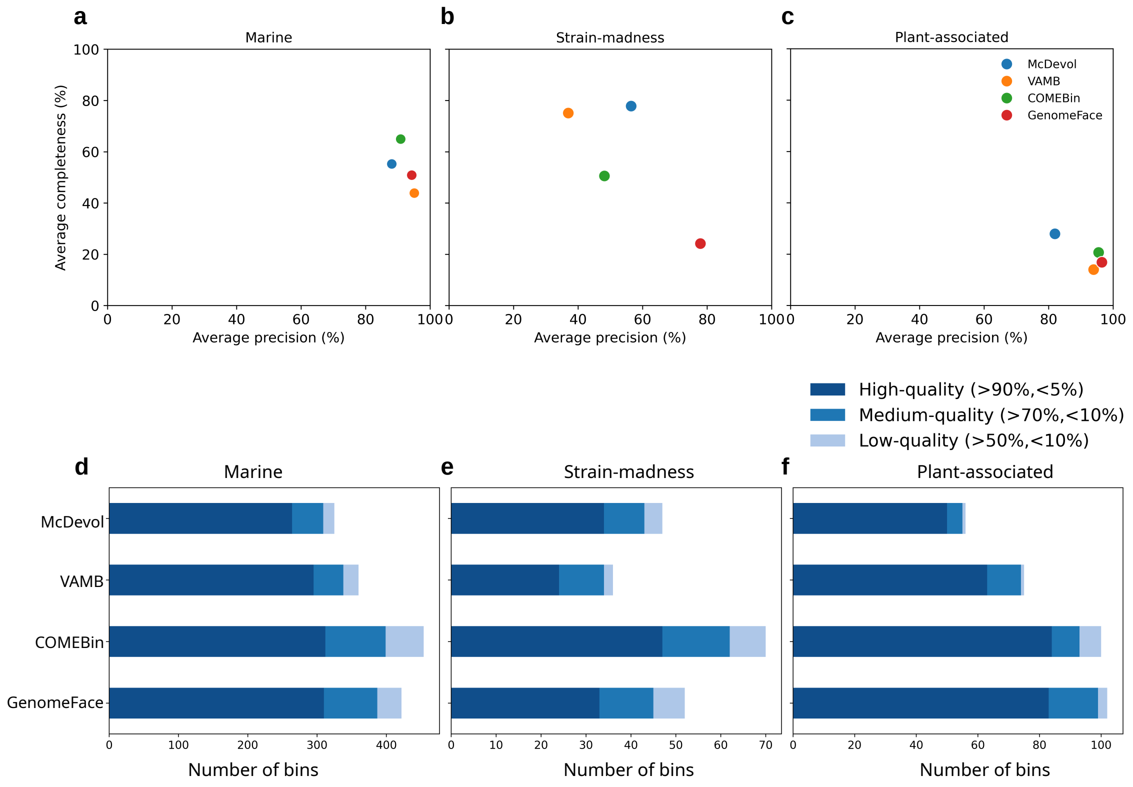


Figure 2: Average purity (bp) and average completeness (bp) of metagenomic bins generated from four deep learning binners for a) marine, b) strain-madness and c) plant-associated datasets. **d-f,** The number of bins in three quality categories generated by McDevol, VAMB, COMEBin and GenomeFace for three CAMI2 datasets.

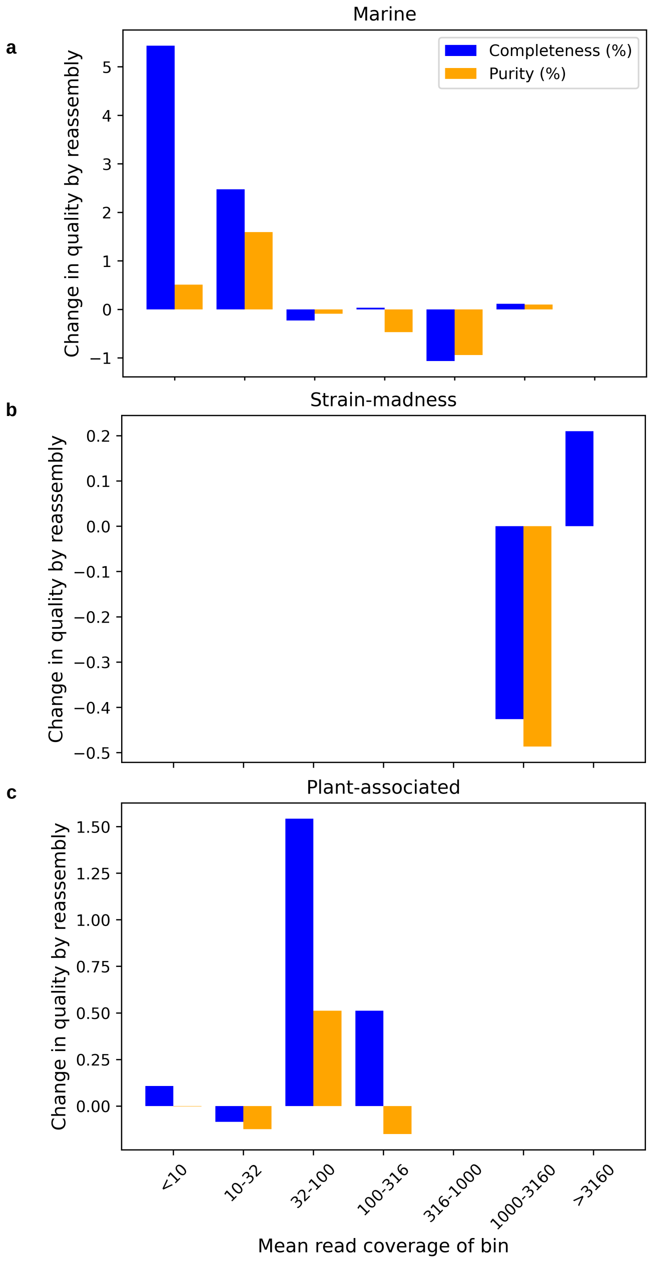


Figure 3: Change in completeness (%) and purity (%) of bins upon reassembly. X axis is the mean of total read coverage of contigs in a bin. Y axis is the difference in values (completeness after reassembly - completeness of bins before reassembly). Positive y axis values indicate improvement in quality values after reassembly. Bin assignment from GenomeFace's results was used for this plot.

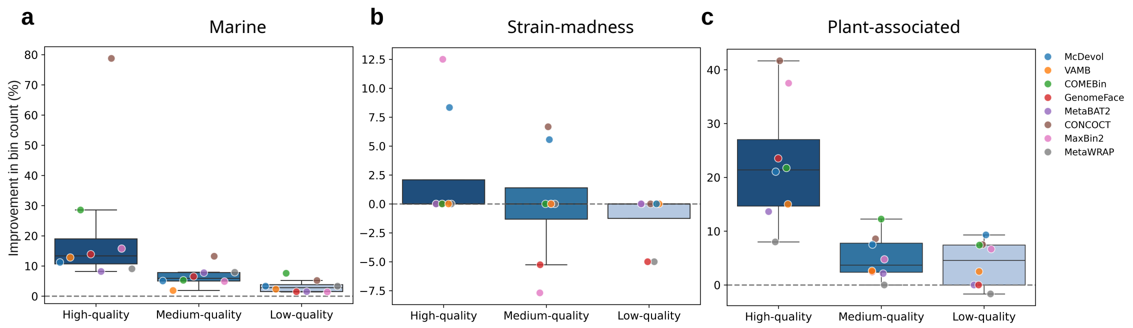


Figure 4: **Evaluation of reassembly after binning.** The percentage increase in the number of bins after reassembly belonging to high, medium and low-quality categories for a) marine, b) strain-madness and c) plant-associated datasets. The percentage increase in the number of bins for each binner is shown as dots.

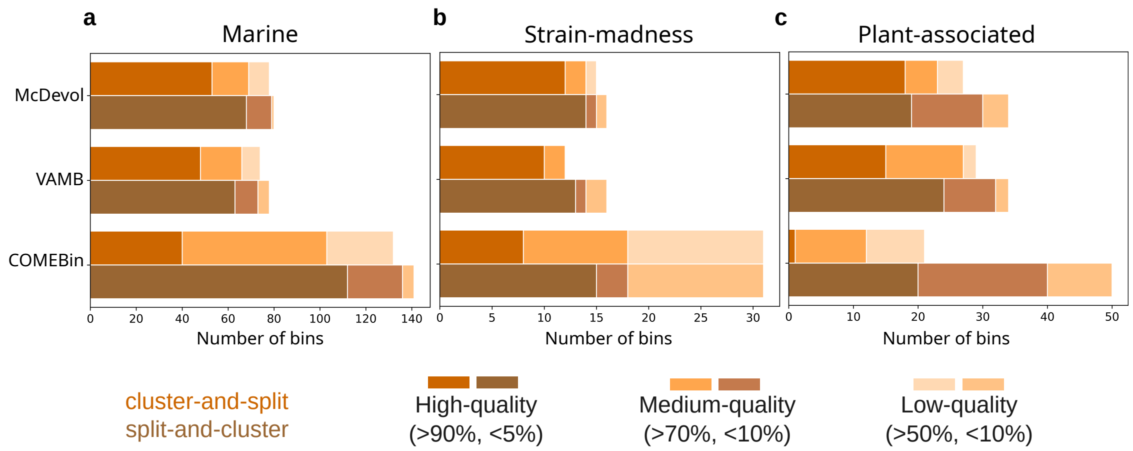


Figure 5: Cluster-and-split *vs* split-and-cluster mode for multi-sample binning. The number of non-redundant high, medium and low-quality bins obtained from two modes of multi-sample binning for a) marine, b) strain-madness and c) plant-associated datasets.

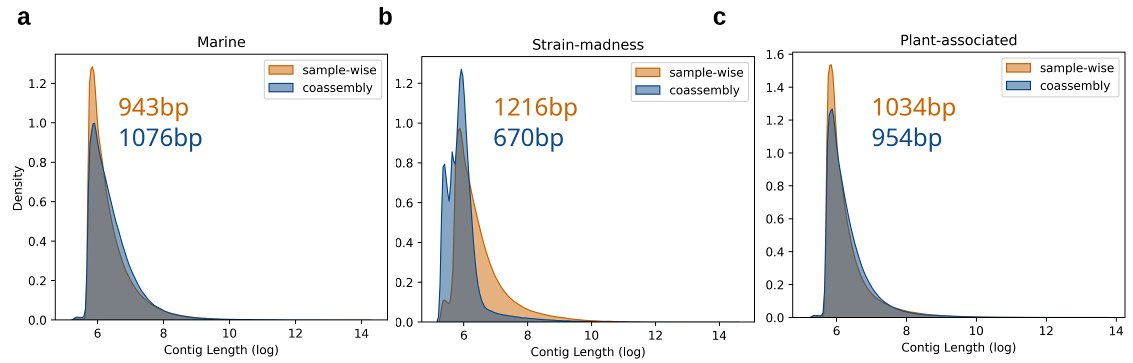


Figure 6: Density distribution of sequence length (base pairs in log scale) of contigs assembled by coassembly (blue) and sample-wise assembly (orange). Numbers written are the mean over contig lengths from different sampling approaches.

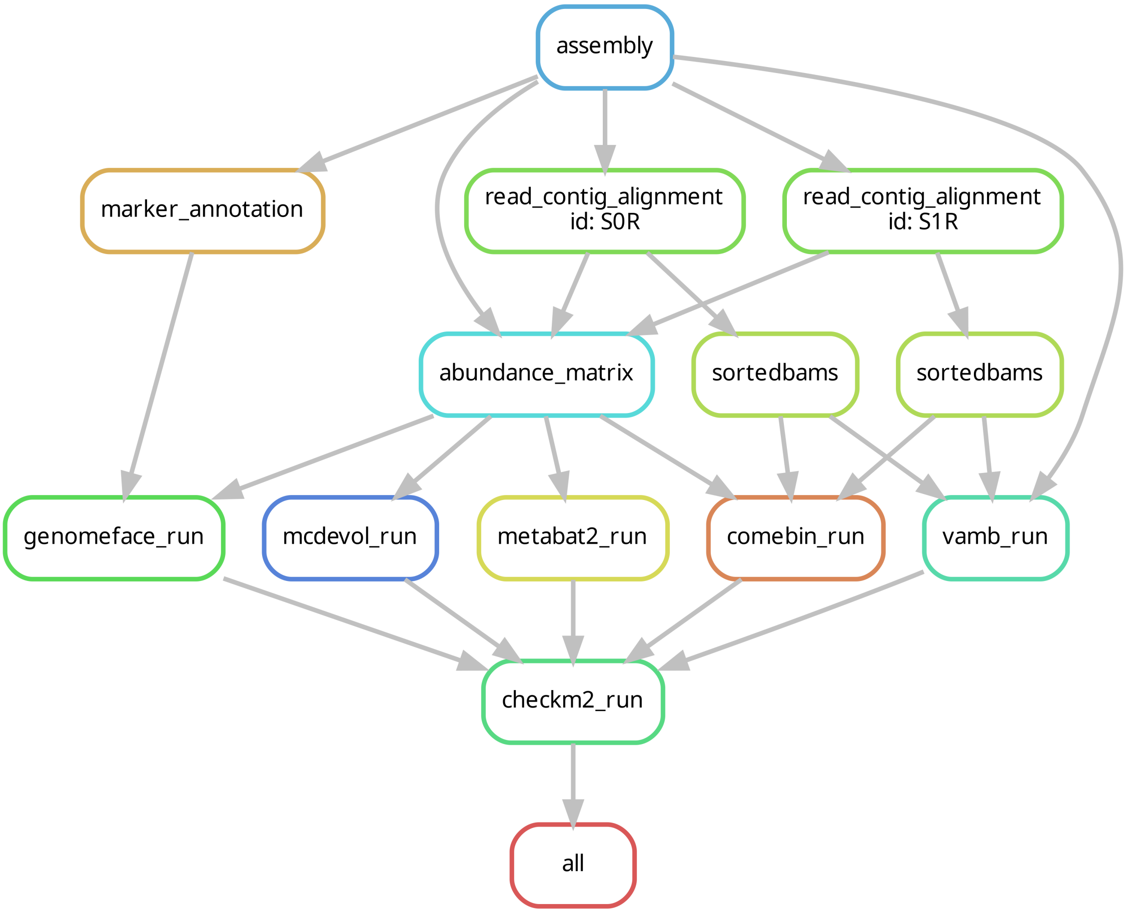


Figure 7: **Workflow for coassembly benchmarking.** It uses MEGAHIT for assembly, Strobealign for read mapping, samtools for alignment sorting, McDevol, VAMB, COMEBin, GenomeFace and MetaBAT2 for binning and CheckM2 for bin evaluation. The arrow indicates that the output of the previous step is used directly as input for the next step.

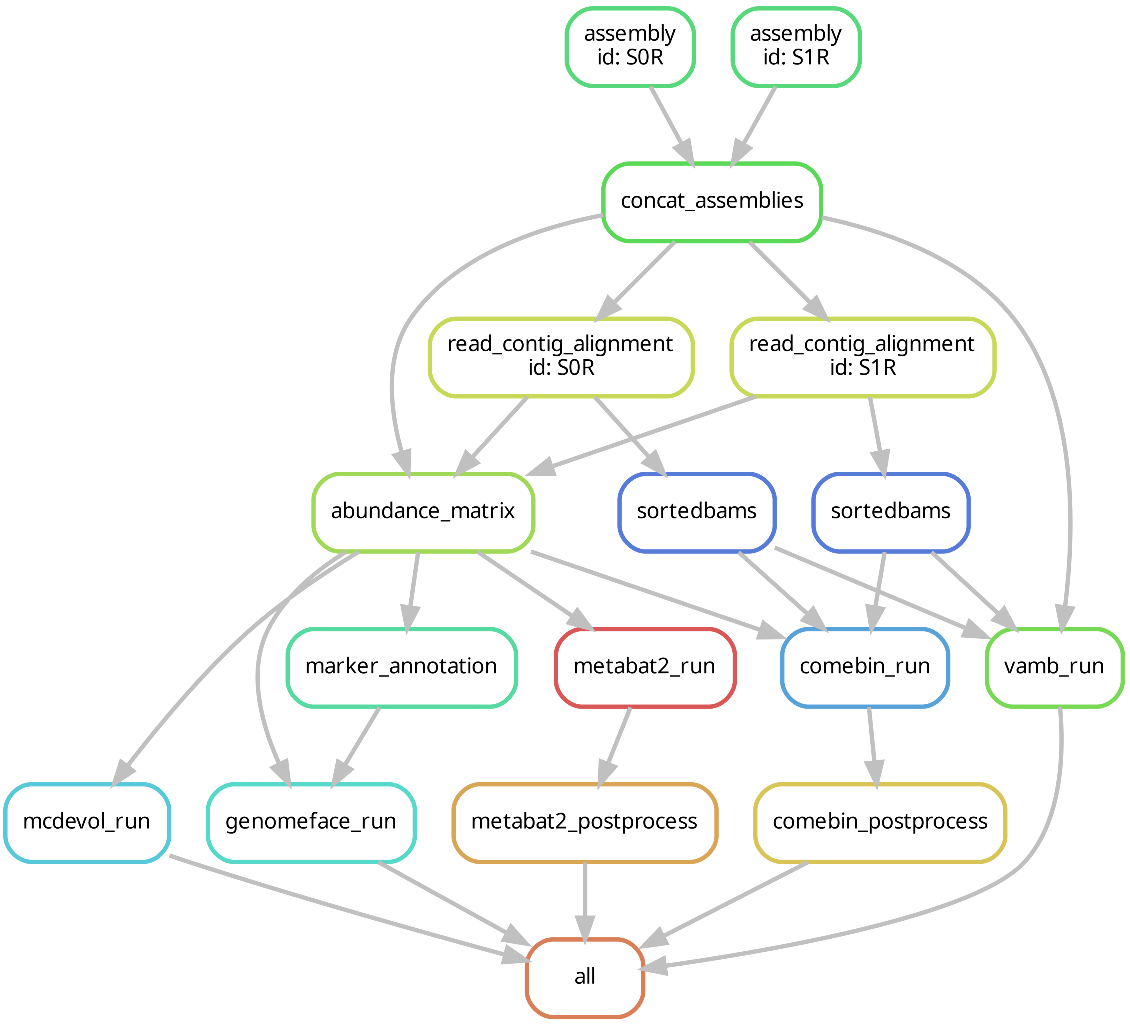


Figure 8: Multi-sample benchmarking workflow that includes assembly, read mapping, alignment, sorting, binning and bin splitting by sample. The tools for each step are the same as those shown in Supplementary Fig.5.

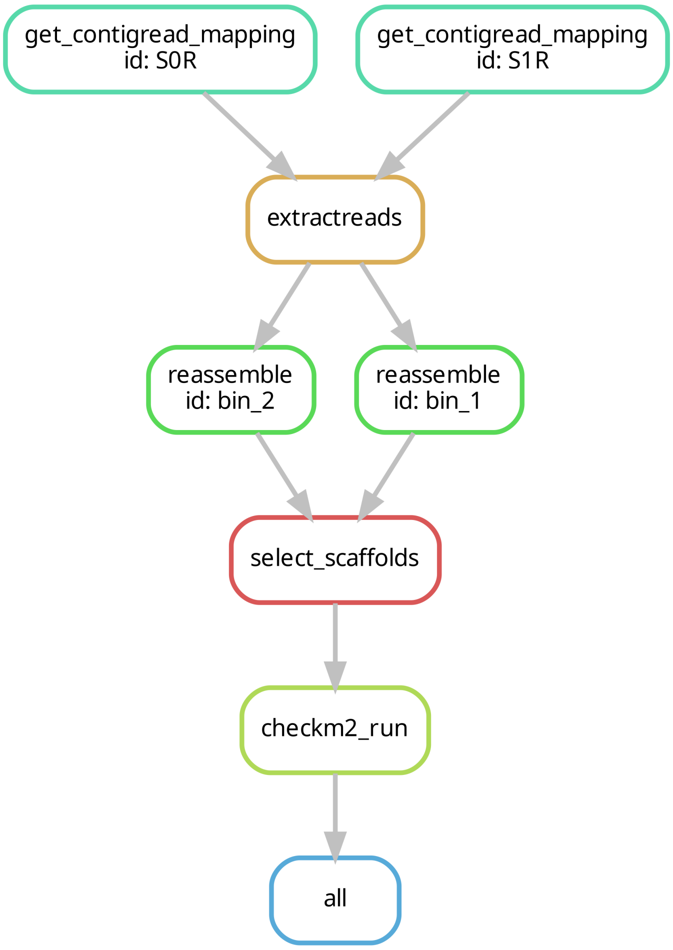


Figure 9: Workflow for post-binning reassembly. This workflow recruits reads mapped to contigs belonging to a bin using our *extractreads* script, reassembles using SPAdes, filters scaffolds to produce final bins using our *convertfasta_multi2single* script, and evaluates using CheckM2. Scripts are available at *https://github.com/soedinglab/binning_benchmarking*.
